# TRIM32 inhibits Venezuelan Equine Encephalitis Virus Infection by targeting a late step in viral entry

**DOI:** 10.1101/2024.06.04.597282

**Authors:** Yifan Xie, Jie Cao, Shuyi Gan, Lingdong Xu, Dongjie Zhang, Suhong Qian, Feng Xu, Qiang Ding, John W. Schoggins, Wenchun Fan

## Abstract

Alphaviruses are mosquito borne RNA viruses that are a reemerging public health threat. Alphaviruses have a broad host range, and can cause diverse disease outcomes like arthritis, and encephalitis. The host ubiquitin proteasome system (UPS) plays critical roles in regulating cellular processes to control the infections with various viruses, including alphaviruses. Previous studies suggest alphaviruses hijack UPS for virus infection, but the molecular mechanisms remain poorly characterized. In addition, whether certain E3 ubiquitin ligases or deubiquitinases act as alphavirus restriction factors remains poorly understood. Here, we employed a cDNA expression screen to identify E3 ubiquitin ligase TRIM32 as a novel intrinsic restriction factor against alphavirus infection, including VEEV-TC83, SINV, and ONNV. Ectopic expression of TRIM32 reduces alphavirus infection, whereas depletion of TRIM32 with CRISPR-Cas9 increases infection. We demonstrate that TRIM32 inhibits alphaviruses through a mechanism that is independent of the TRIM32-STING-IFN axis. Combining reverse genetics and biochemical assays, we found that TRIM32 interferes with genome translation after membrane fusion, prior to replication of the incoming viral genome. Furthermore, our data indicate that the monoubiquitination of TRIM32 is important for its antiviral activity. Notably, we also show two TRIM32 pathogenic mutants R394H and D487N, related to Limb-girdle muscular dystrophy (LGMD), have a loss of antiviral activity against VEEV-TC83. Collectively, these results reveal that TRIM32 acts as a novel intrinsic restriction factor suppressing alphavirus infection and provides insights into the interaction between alphaviruses and the host UPS.

**Author summary:** Due to climate change, wildlife habitat loss, and human activities, alphavirus infections are a growing threat to public health. The host UPS has critical role in virus-host interaction, but how the UPS impact alphavirus infection is not completely understood. In this study, we found that the E3 ubiquitin ligase TRIM32 inhibits diverse alphaviruses in multiple cell types. Mechanistically, TRIM32 impairs primary translation of incoming viral genome in a manner that depends on. monoubiquitination of TRIM32. Additionally, disease-associated alleles of TRIM32 have a loss-of-function with respect to viral inhibition. Together, these findings uncover a novel biological function of TRIM32 in regulating alphavirus infection and provide important insights into the interplay between alphaviruses and the host USP.

## Introduction

Alphaviruses are mosquito transmitted positive-sense single-stranded RNA viruses that have a broad host range, including human, horses, birds, and rodents [1]. Based on the genetic relatedness and disease manifestation, alphaviruses are classified into two groups, arthritogenic alphaviruses and encephalitic alphaviruses. The former include chikungunya virus (CHIKV), O’ynong-nyong virus (ONNV), Ross river virus (RRV), and Mayaro virus (MAYV), and the latter include Venezuelan equine encephalitis virus (VEEV), eastern and western equine encephalitis virus (EEEV, WEEV) [2,3]. To date, there is no approved vaccine or specific antiviral agent for treating encephalitic alphavirus infection. Due to climate change, human activities, and habitat loss of wild animals, alphavirus infections have been identified globally and pose potential threats to humans worldwide [4,5]. Accordingly, understand how these viruses interact with their hosts at the molecular level is an important area of research.

Both host factors and restriction factors are important for governing virus infection. Identification and characterization of these factors offers valuable insight into viral pathogenesis and potential strategies to develop of antiviral therapeutics. Genetic screens have been used to identify numerous host factors that are important for alphavirus infection. These include cellular receptors for alphavirus entry, such as MXRA8 for CHIKV and ONNV [6], LDLRAD3 for VEEV [7], and VLDLR and ApoER2 for Semliki forest virus (SFV) [8]. Host factors that are required for alphavirus replication have also been identified, including G3BP1/2, FXR1/2, FMR [9], SEC61A, VCP [10], and FHL1 [11]. Several interferon-stimulated gene (ISG) products, including ZAP, IFIT1, IFIT3, ISG20, viperin, and several poly(ADP-ribose) polymerases, have been implicated as anti-alphavirus restriction factors. [12–16]. In addition to ISGs, basally expressed proteins such as DDX56 [17] and DDX39A [18] have been identified as restriction factors that inhibit alphaviruses.

Previous studies found that the host ubiquitin proteasome system (UPS) plays a crucial role in alphaviruses infection. Specifically, the post-entry stages of alphaviruses can be disrupted by UPS inhibitors, such as proteasome inhibitors and deubiquitinating enzyme inhibitors [19–21]. To date, only a few E3 ubiquitin ligases (E3) and deubiquitinases (DUB) have been identified as host factor or restriction factor for alphaviruses. TRIM25, a well-studied E3 that participates in host antiviral innate immunity [22], also acts as a co-factor of zinc-finger antiviral protein (ZAP) to promote ZAP-mediated inhibition of alphaviruses [23–25]. In mosquito cells, an E3 ubiquitin ligase scaffolding protein, AcCullin3 was identified as pro-viral factor of CHIKV replication [26]. The membrane associated ring-CH-type finger 8 (MARCH8) is a broad-spectrum antiviral E3, which can also restrict alphavirus infection by targeting viral envelope protein E2 for degradation [27,28].

In this study, we used a cDNA expression strategy to screen 118 human E3 ligases for antiviral effects on VEEV-TC83-GFP infection. TRIM32 was identified as a novel restriction factor that inhibits multiple alphaviruses including VEEV-TC83, SINV, and ONNV. TRIM32 is a RING type E3 ubiquitin ligase [29] that has reported roles in influenza A virus infection, tumorigenesis, and microRNA processing [29–33]. TRIM32 is also reported to be involved in regulation of the host antiviral immune response. TRIM32-mediates the K63-linked ubiquitination of MITA/STING to promote STING-mediated interferon production and cellular antiviral responses [34]. TRIM32 also targets TRIF for TAX1BP1-mediated selective autophagic degradation to suppress TLR3/4-mediated immune responses [35]. TRIM32 can also directly regulate virus infection. It has been reported that TRIM32 regulates the biological activity of the HIV-1 Tat protein [36] and directly targets PB1 protein of Influenza A virus (IAV) for proteasome degradation to restrict viral replication [37].

To dissect the underlying mechanisms of TRIM32-mediated inhibition of alphavirus, we used VEEV-TC83 as a model. By combining reverse genetic and biochemical assays, we show that TRIM32 limits the translation of incoming viral genome. Proximity-labeling proteomic approaches and intracellular co-localization analysis suggest TRIM32 associates with endosome-lysosome compartments, where alphaviruses undergo membrane fusion and uncoating to initiate cytosolic entry [38]. Furthermore, we found that two pathogenic mutants of TRIM32 associated with Limb-girdle muscular dystrophy type 2H (LGMD2H), R394H and D487N, lack antiviral activity against VEEV-TC83. Together, our findings highlight TRIM32 as cell intrinsic restriction factor that inhibits alphavirus infection.

## Results

### TRIM32 inhibits VEEV infection

To identify E3 ubiquitin ligases that modulate alphavirus infection, we screened 118 human RING-type E3 ubiquitin ligases for effects on VEEV-TC83-GFP in HeLa cells using methods described in our previous study [39]. Briefly, HeLa cells were transduced with E3-expressing lentiviruses and were then infected with VEEV-TC83-GFP at MOI of 0.1 for 24 hours (hrs), after which virus infectivity was quantified by flow cytometry. We found that VEEV-TC83-GFP infectivity was strikingly reduced in TRIM32 expressing cells, and enhanced by TRIM33 (Fig 1A). Next, we established HeLa stable cells expressing TRIM8, TRIM25, TRIM26, TRIM32, and TRIM33 to evaluate their effects on VEEV infection. Again, TRIM32 showed antiviral activity, while TRIM33 showed pro-viral activity towards VEEV (Fig 1B). The other three TRIM proteins did not influence VEEV infection, even though they are reported to positively regulate host antiviral responses [40]. We further investigated the effects of TRIM32 on other RNA viruses, coxsackievirus B3 (CVB3), Zika virus (ZIKV), and vesicular stomatitis virus (VSV). In contrast to VEEV, these viruses were not restricted by TRIM32 expression (S1A Fig). Based on these screening results, we chose TRIM32 for additional mechanistic studies.

**Fig 1.**
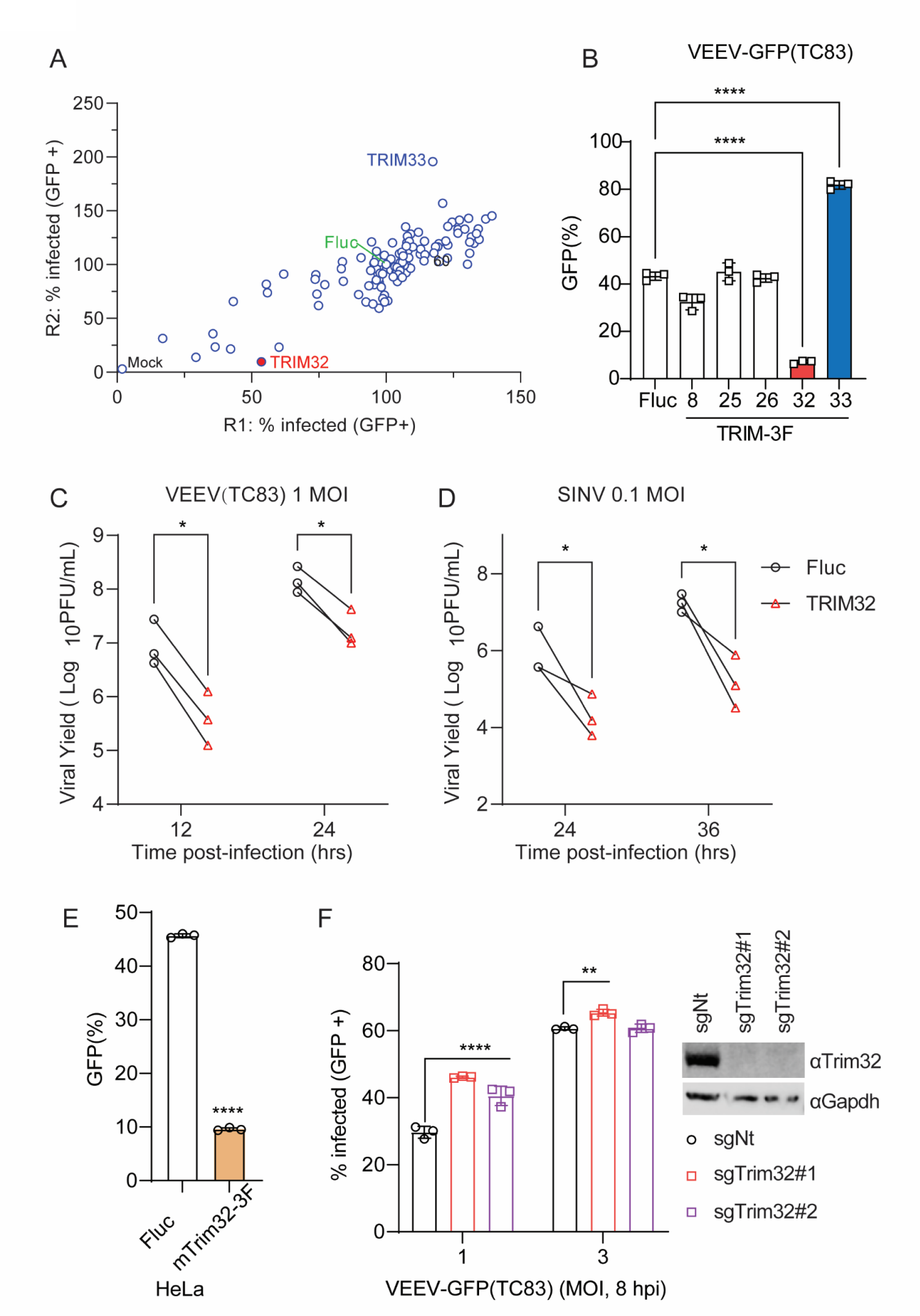
Identification of TRIM32 as an antiviral effector against alphaviruses. **A**. HeLa cells transduced with lentivirus co-expressing RFP and 118 different human RING type E3 ubiquitin ligases were infected with VEEV-TC83-GFP. Virus infectivity was determined as the percentage of GFP-positive cells within the RFP-positive gate using flow cytometry. Infection values were normalized to the empty vector control. **B**. HeLa cells expressing the indicated E3 or Fluc control were infected with VEEV-TC83-GFP at MOI of 0.1, and virus infectivity was measured at 24hpi by flow cytometry. **C** and **D**. HeLa-Fluc or HeLa-TRIM32 cells were infected with VEEV (**C**) or SINV (**D**) at the indicated MOI. The progeny virions in culture medium were collected at the indicated time points post-infection. and quantified by plaque assay. **E**. HeLa cells expressing mouse Trim32 or empty plasmid control infected with VEEV-TC83-GFP at MOI of 0.1 for 24h. Viral infectivity was quantified by flow cytometry. **F**. Left: C2C12 cells transduced withTrim32 targeting sgRNA or non-targeting sgRNA were infected VEEV-TC83-GFP at the indicated MOI for 8 h, and viral infectivity was determined by flow cytometry. Right: Western blot analysis of Trim32 or housekeeping Gapdh on the indicated C2C12 cells. Data represent averages of independent biological replicates and are presented as means ± SD (n=3). Statistical significance was determined by unpaired students’ t-test (*P<0.05, **P<0.01, ****P<0.0001).

To determine whether TRIM32 retained antiviral activity against non-reporter alphaviruses, HeLa cells stable expressing Fluc or TRIM32 bearing a 3X-FLAG tag (TRIM32-3F) were infected with VEEV-TC83 and SINV at MOI of 1 and 0.1, respectively. The cell culture supernatants were collected after multiple viral life cycles, and the production of progeny virions was determined by plaque assay. Both VEEV-TC83 and SINV production significantly decreased in TRIM32-expressing cells compared to control cells (Fig 1C and 1D). Because rodent animals play a role in alphavirus transmission, we speculated that mouse Trim32 may also influence alphavirus infection. Indeed, we found that Trim32 expression restricted VEEV-TC83-GFP infection significantly (Fig 1E). To complement the ectopic expression effects of TRIM32, we used CRISPR-Cas9 editing to silence Trim32 in C2C12 cells, which highly express Trim32. We observed a modest rescue of infection in the Trim32-depleted cells, suggesting that endogenous TRIM32 may contribute to cell intrinsic suppression of VEEV (Fig 1F). Together, these data demonstrate that TRIM32 from multiple species may function as an alphavirus restriction factor.

### TRIM32 mediated inhibition of VEEV is independent of the TRIM32-STING-IFN axis

Previous studies have reported that TRIM32 is a positive regulator of STING-mediated interferon production and cellular innate immune responses [34]. Therefore, we sought to determine if TRIM32 inhibits alphaviruses through regulation of STING or by other mechanisms. We used CRISPR-Cas9 editing to silence STING in HeLa cells and then test the effects of TRIM32 on alphavirus infection in the absence of STING [41]. Notably, TRIM32 retained anti-VEEV activity even when STING was depleted in HeLa cells (Fig 2B). Similar results were observed for U2-OS cells (S1B Fig). In addition, we deleted TBK1, a kinase that phosphorylates multiple regulators in of antiviral immunity [42], in TRIM32 expressing cells. Again, VEEV-TC83-GFP infection was strikingly reduced by TRIM32 expression in the absence of TBK1 (Fig 2C). Similar antiviral effects of TRIM32 were observed in Huh7 cells, which lack detectable levels of STING (Fig 2D) [41]. Finally, silencing of two key transcriptional regulators of the interferon signaling pathway, STAT1 and IRF9, also did not abolish TRIM32-mediated inhibition of VEEV-TC83-GFP, but did abrogate interferon-mediated inhibition of the virus (Fig 2E, 2F). Collectively, our findings demonstrate that TRIM32 inhibits alphavirus independently of canonical of interferon-mediated antiviral pathways.

**Fig 2.**
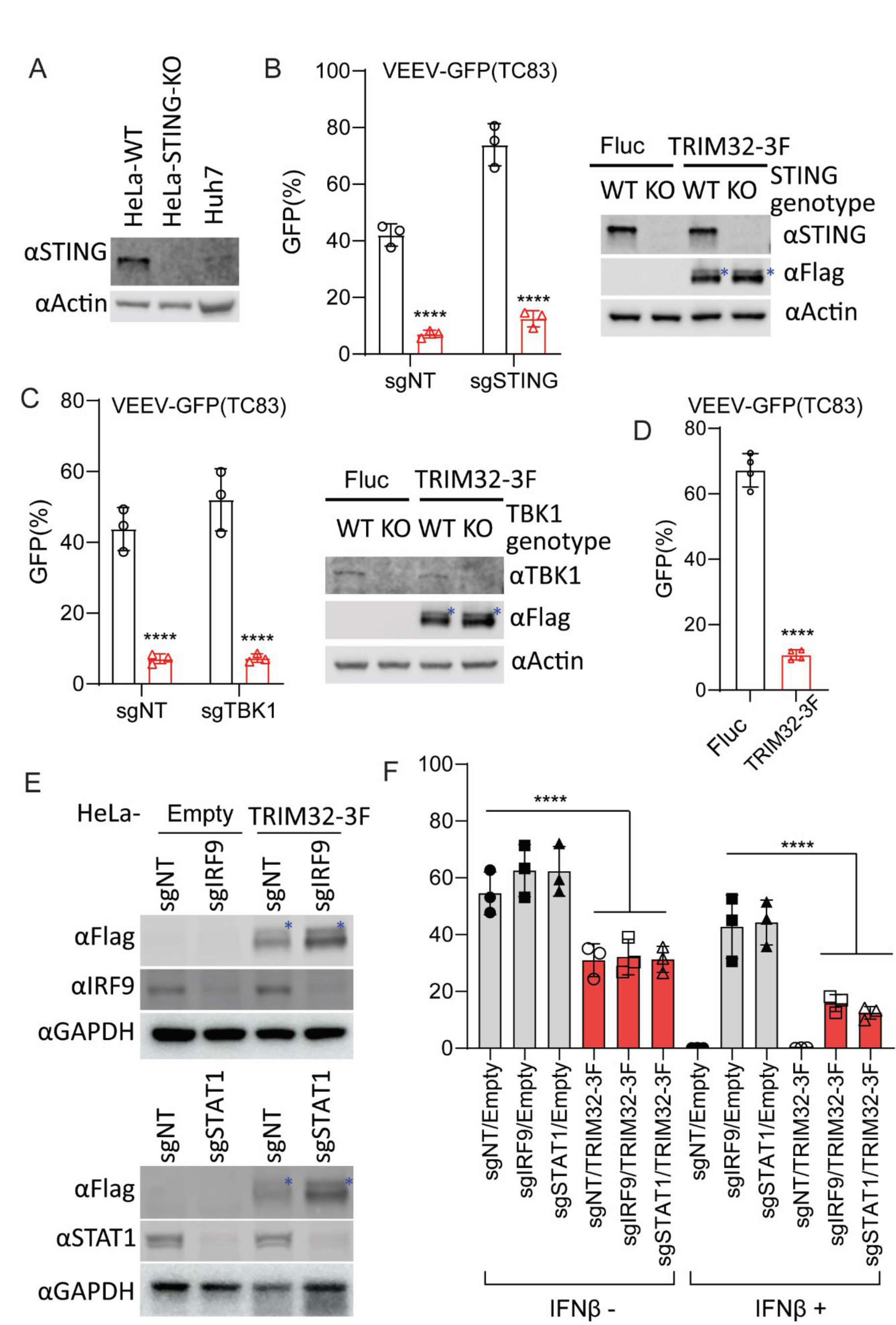
TRIM32-mediated alphavirus inhibition is independent of the TRIM32-STING-IFN axis. **A**. Western blot analysis of STING protein expression from the indicated cells. **B**. HeLa-sgSTING or HeLa-sgNT with or without TRIM32 expression were infected VEEV-TC83-GFP at MOI of .01 or 1 for 24h, and viral infectivity was quantified by flow cytometry. The asterisks denote the monoubiquitinated TRIM32. **C**. HeLa-sgTBK1 or HeLa-sgNT with or without TRIM32 expression were infected VEEV-TC83-GFP at MOI of .01 1 for 24h, and viral infectivity was quantified by flow cytometry. The asterisks denote the monoubiquitinated TRIM32. **D**. Huh7 cells expressing mouse TRIM32 were infected with VEEV-TC83-GFP at 0.1 MOI for 12h, virus infectivity was quantified by flow cytometry. **E**. Western blot analysis of IRF9 and STAT1 from the indicated cells. The asterisks denote the monoubiquitinated TRIM32. **F**. The indicated cells were pretreated with IFNβ at a concentration of 100U for 16h, then infected with VEEV-TC83-GFP at 0.1 MOI for 24h, and viral infectivity was determined by flow cytometry. Data represent averages of independent biological replicates and are presented as means ± SD (n=2-4). In Statistical significance was determined by unpaired students’ t-test (****P<0.0001).

### TRIM32 does not influence the post-entry stages of VEEV

To determine which step in the viral replication cycle is targeted by TRIM32, we first performed a post-entry infection assay. We transfected VEEV-TC83-GFP infectious viral RNA into HeLa-TRIM32 and HeLa-Fluc cells to measure virus production when entry is bypassed. The progeny virions in supernatants were collected at multiple time points up to 12 hrs and reinfected on to naïve Huh7 cells. Infectivity of the newly produced viruses was then quantified by flow cytometry. By this method, no difference was observed between HeLa-TRIM32 and HeLa-Fluc cells (Fig 3A). Similar results were obtained by transfection of non-reporter VEEV-TC83 viral RNA, followed by quantification of viral yield by plaque assay (Fig 3B). At a later time point of 18 hrs, the viral yield was reduced in HeLa-TRIM32 cells relative to control cells (Fig 3C). These data indicate that TRIM32 has no effect on post-entry stages of VEEV-TC83 infection when viral replication is launched from transfected RNA.

**Fig 3.**
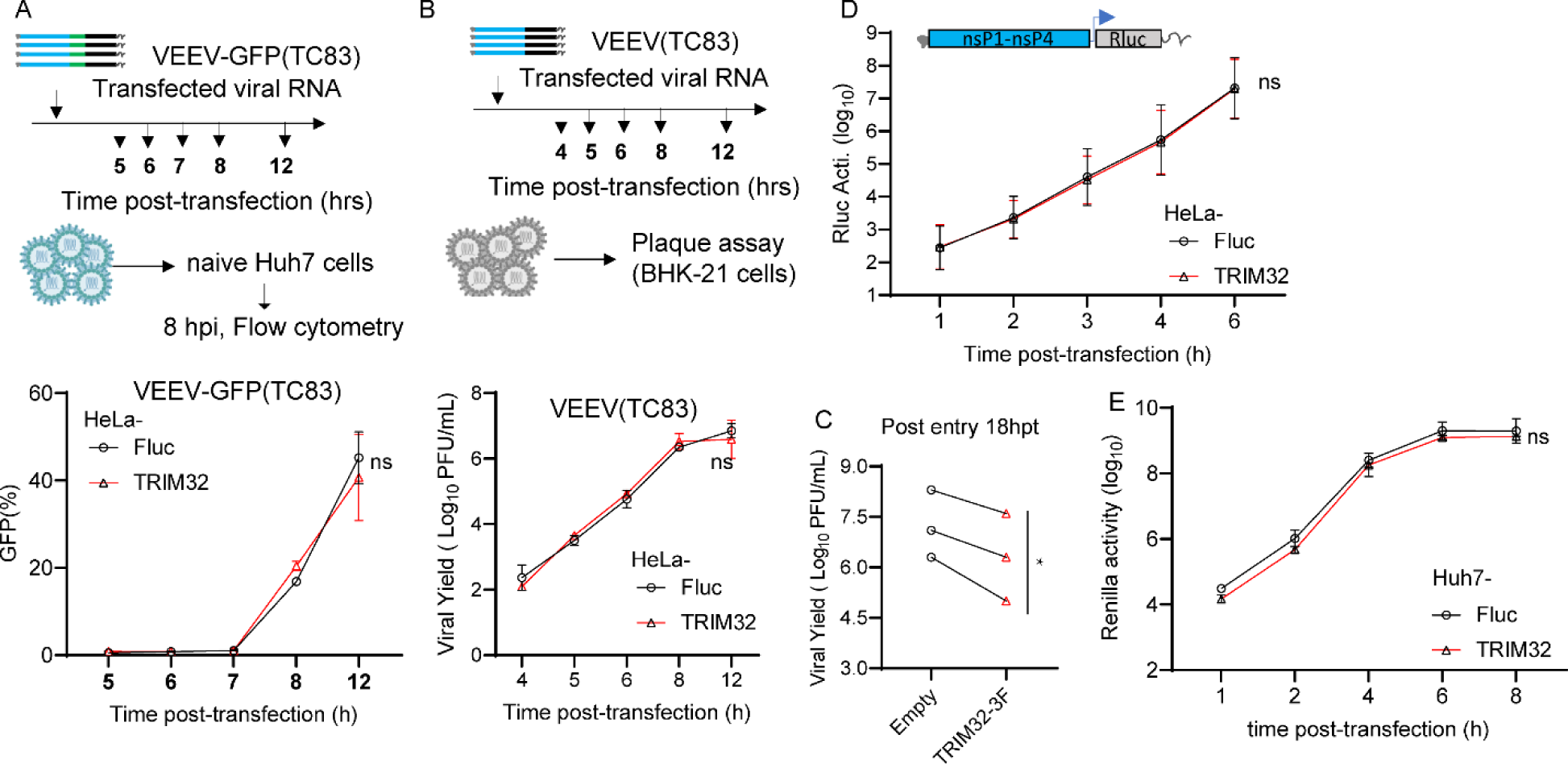
TRIM32 does not influence post-entry stages of the VEEV infection cycle. **A**. Upper: schematic of post-entry assay by transfecting VEEV-TC83-GFP infectious RNA on HeLa-empty or HeLa-TRIM32 cells. Bottom: the supernatants were harvested at the indicated time points post-transfection and used to infect naïve HeLa cells. Infectivity of progeny virus was determined by flow cytometry. **B**. Upper schematic of post-entry assay by transfecting VEEV-TC83 infectious RNA on HeLa-Fluc or HeLa-TRIM32 cells. Bottom: the supernatants were collected at the indicated time points post-transfection. The progeny virus in supernatants were determined by plaque assay on BHK-21 cells. C. HeLa-Fluc or HeLa-TRIM32 cells were transfected with VEEV-TC83 infectious RNA for 18 hrs, the supernatants were collected and the progeny virus in supernatants were determined by plaque assay on BHK-21 cells. **D**. HeLa-Empty or HeLa-TRIM32 cells were transfected with VEEV-TC83-Renilla luciferase (Rluc) replicon RNA. The cells were harvested at indicated time points, and Rluc activity was quantified. **E**. VEEV-TC83-Renilla luciferase (Rluc) replicon assay on Huh7-Empty or Huh7-TRIM32. Three independent biological replicates were carried out for each experiments. Statistical significance was determined by 2-way ANOVA with Sidak’s multiple comparisons test for A, B, D, E, ns, no significant. A paired t-test was used for C (*P<0.05).

To confirm this, we carried out a VEEV replicon assay. VEEV-TC83 replicon RNA expressing Renilla luciferase was transfected into HeLa-TRIM32 or HeLa-Fluc cells, and cells were collected at multiple time points post-transfection to measure Renilla luciferase activity. The results demonstrate that the VEEV-TC83 replicon produced similar levels of Renilla luciferase in both TRIM32 and Fluc expressing cells, suggesting TRIM32 does not influence translation of transfected viral genomes and subsequent replication (Fig 3D). Similar results were observed in Huh7 cells (Fig 3E). Together, these data suggest TRIM32 does not impair the post-entry steps of VEEV infection.

### TRIM32 does not inhibit VEEV binding, entry, or membrane fusion

We next sought to investigate effects of TRIM32 on the early stages of VEEV infection. Starting with a binding and entry assay, we did not observe statistically significant changes in viral binding or entry of VEEV-TC83 when comparing HeLa cells expressing TRIM32 and Fluc (Fig 4A). However, a modest decrease (< 2-fold) of viral binding was observed in HeLa-TRIM32 cells. To verify this observation with an assay that might offer more sensitivity, we established a VEEV that contains a Nano-luciferase (NLuc) fusion at the N-terminus of capsid, hereafter termed VEEV-NLuc/Cap, as a tool to study VEEV binding (S2A Fig). Because the NLuc is a part of the VEEV-NLuc/Cap virion (S2B Fig), we can evaluate virus binding efficiency by measuring NLuc activity. To ensure this modification has no influence on receptor usage of LDLRAD3, we established HeLa-sgLDLRAD3 cells by using a human LDLRAD3 specific sgRNA for CRISPR-mediated gene silencing [7], followed by virus infectivity determination. As shown in Figure S2C and S2D, both VEEV-GFP and VEEV-NLuc/Cap infection was reduced on HeLa-sgLDLRAD3 cells, but not ONNV and SINV. Finally, our data showed that there is no significant difference in NLuc activity after cold-binding of VEEV-NLuc/Cap to TRIM32 and Fluc expressing cells (Fig 4B and 4C).

**Fig 4.**
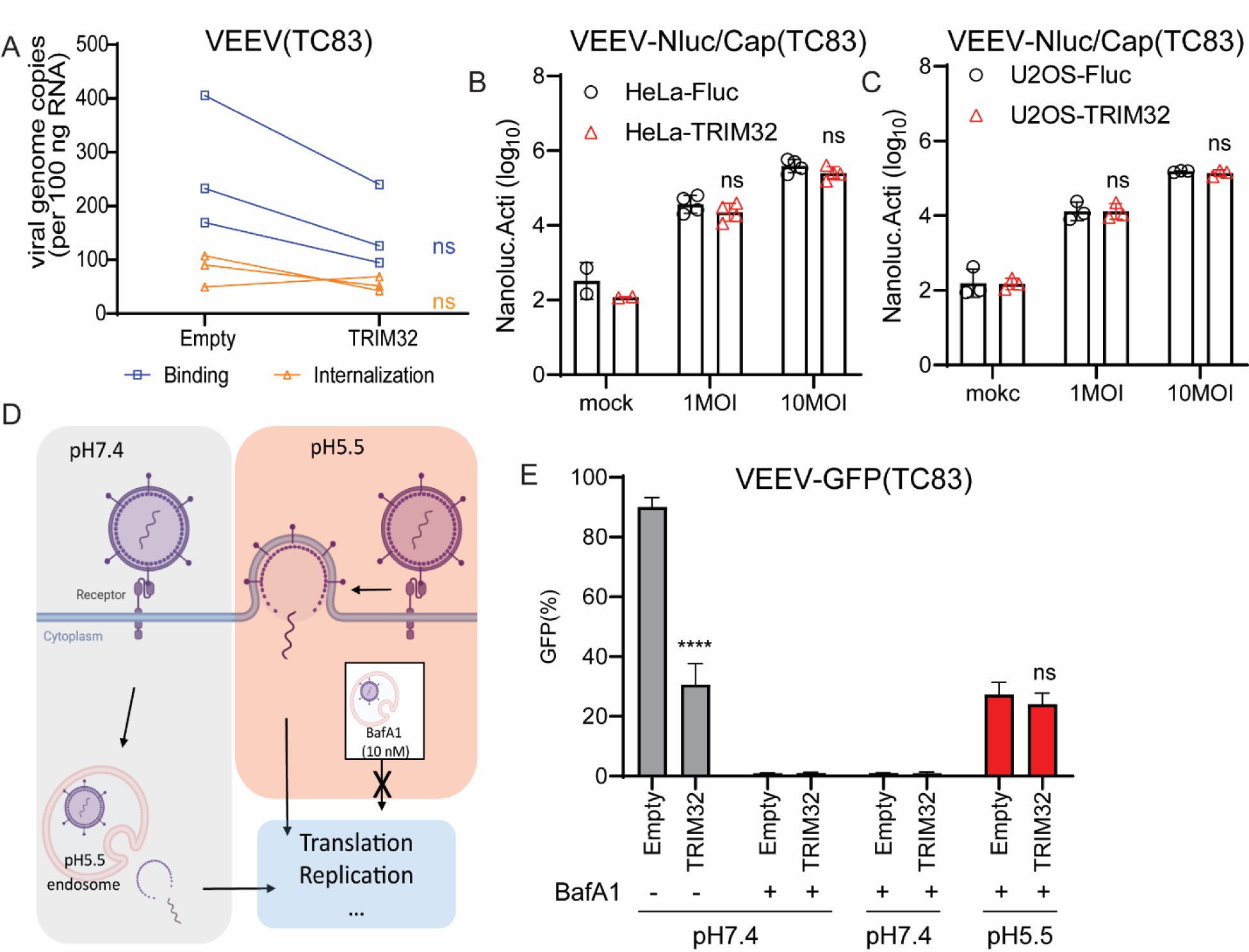
TRIM32 does not impair viral binding, entry, and membrane fusion. **A.** HeLa-Empty or HeLa-TRIM32 cells were infected with 50 MOI of VEEV-TC83 at 4°C for 1.5h, and bound virions were quantified by real-time qPCR assay. **B**. HeLa-Empty or HeLa-TRIM32 cells were infected with VEEV-NLuc/Cap at 4°C for 1.5h, then the cells were washed and collected to determine Nanoluc activity. **C**. U2OS-Empty or U2OS-TRIM32 cells were infected with VEEV-NLuc/Cap at 4°C for 1.5h, then the cells were washed and collected for to determine Nanoluc activity. **D**. Schematic of acid bypass assay. **E.** Acid bypass assay of VEEV-TC83-GFP on HeLa-Empty or HeLa-TRIM32 cells. Viral infectivity was quantified by flow cytometry. Data represent averages of independent biological replicates and are presented as means ± SD (n=3). Statistical significance was determined by unpaired students’ t-test for A, B, C, and G (****P<0.0001). For D and E, statistical significance was determined by 2-way ANOVA with Sidak’s multiple comparisons test (****P<0.0001).

Next, we asked whether TRIM32 targets viral membrane fusion by performing an acid bypass assay as previously described (Fig 4D) [43]. Briefly, HeLa-TRIM32 or HeLa-Fluc cells were infected with VEEV-TC83-GFP at MOI of 10 at 4°C for 1.5h. The unbound virions were washed using cold PBS, then the cells were incubated in PBS at pH7.4 or pH5.5 at 4°C for 10 minutes, with the more acidic pH promoting to viral membrane fusion on the plasma membrane. After neutral or acidic PBS treatment, PBS was replaced with pre-warmed medium containing 10nM bafilomycin A1 (BafA1) to block endosomal escape and incubated at 37°C for 12h. Viral infectivity was then quantified by using flow cytometry. We found that VEEV-TC83-GFP displayed similar infectivity under acidic conditions that bypass endosome-lysosome mediated membrane fusion in HeLa-TRIM32 and HeLa-Fluc cells (Fig 4E). Together, these findings suggest that TRIM32 does not obviously affect VEEV early steps of viral binding or membrane fusion.

### TRIM32 impairs the cytosolic entry of VEEV after endosomal membrane fusion prior to genome translation

To further investigate the potential mechanisms of TRIM32-mediated VEEV inhibition, we attempted to identify viral and cellular proteins that interact with TRIM32 during viral infection. To do so, we applied a TurboID-mediated proximity labeling proteomics approach (Fig 5A) [44]. We first established HeLa stable cells expressing TRIM32/TurboID/HA or TurboID/HA and confirmed HeLa-TRIM32/TurboID/HA still exhibits inhibition of VEEV-TC83 (Fig 5B). Next, the cells were mock-infected or infected with VEEV-TC83-GFP at MOI of 25 in the presence of 500μM biotin for 6 hrs. Proximity labeling proteomic analysis was then performed as described previously [44]. After pulldown of candidate proteins with streptavidin beads and proteomic analysis, both TurboID.HA and TRIM32/TurboID/HA were shown to successfully target certain proteins for biotinylation (S3A Fig). Surprisingly, we found that both VEEV structural proteins and nonstructural proteins were enriched by TRIM32/TurboID/HA (Fig 5C). Compared to TurboID/HA, there were 297 cellular proteins enriched in TRIM32/TurboID/HA samples, and 38 cellular proteins shared in TRIM32/TurboID/HA groups that were mock-infected or infected with VEEV-TC83 (S3B Fig).

**Fig 5.**
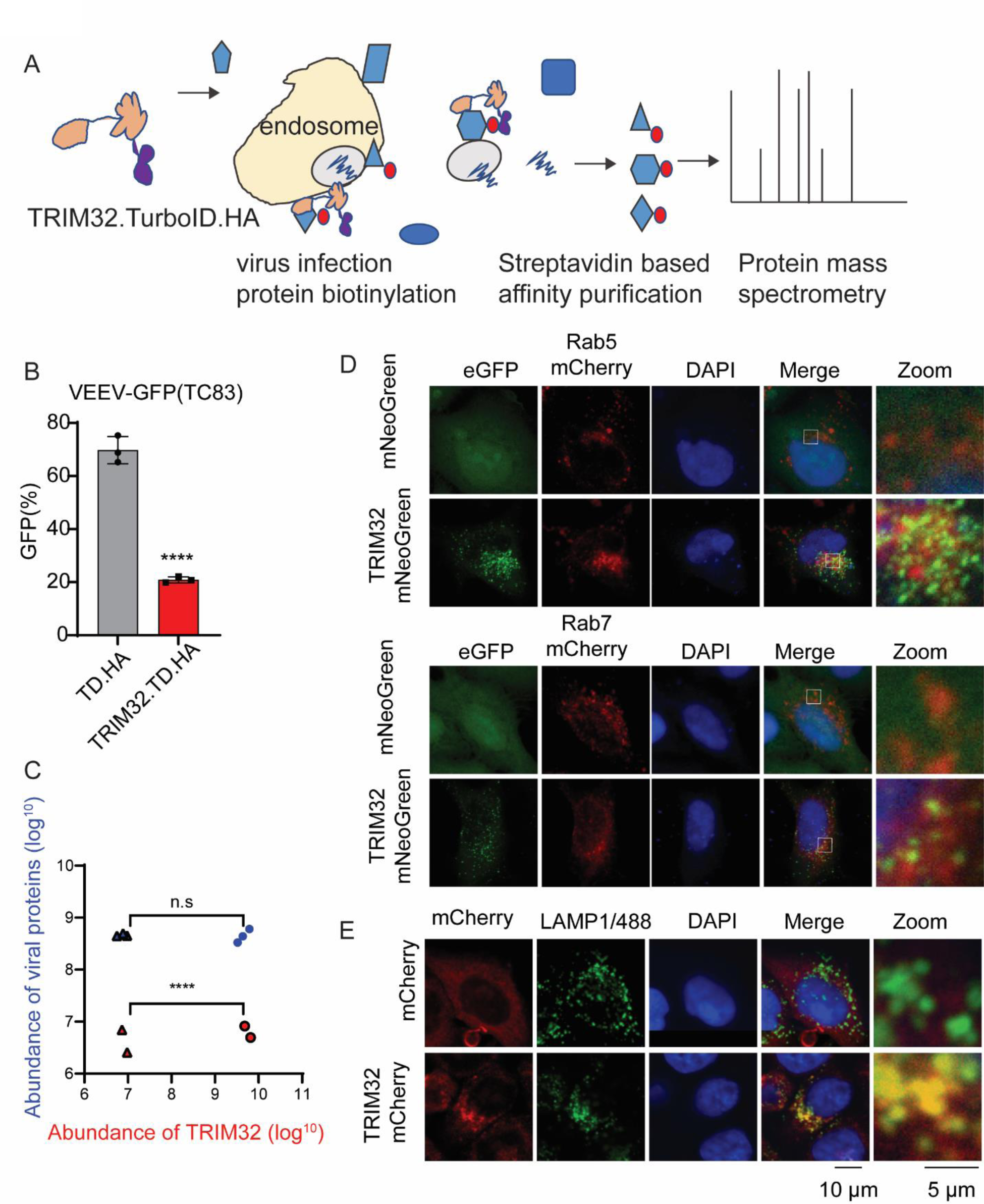
Proximity-labeling proteomic analysis suggests TRIM32 associates with the endosome-lysosome compartment. **A.** Schematic of proximity labeling proteomic approach for identification of cellular proteins and viral proteins of VEEV that interact with TRIM32. **B.** HeLa-TurboID.HA or HeLa-TRIM32.TurboID.HA cells were infected with VEEV-TC83-GFP at 0.1 MOI for 12h, virus infectivity was quantified by flow cytometry. **C.** The protein abundance of TRIM32 and viral structural and nonstructural polyproteins from the indicated groups. Triangle, HeLa-TurboID.HA infected with (blue) or without (red) infection. Circle, HeLa-TRIM32.TurboID.HA infected with (blue) or without (red) infection. **D-E.** Confocal images of the intracellular localization of TRIM32 and the indicated endosome or lysosome markers. Data represent averages of independent biological replicates and are presented as means ± SD (n=2-3). Statistical significance was determined by unpaired students’ t-test for A and C (****P<0.0001).

We analyzed the functions of the cellular proteins enriched by TRIM32 by using PANTHER [45], and found groups of proteins associated with endosomes or the endosome membrane, which are relevant to VEEV infection (S3C Fig). Those proteins included RAB5B, WDR81, and PIK3C3, which are known endosomal proteins (S3D Fig). To validate TRIM32 associates with endosome, we used co-localization analysis by microscopy. Our confocal images show that overexpressed TRIM32-mNeoGreen co-localizes with Rab7-mCherry, and TRIM32-mCherry (Fig 5D) co-localizes to endogenous LAMP1 (Fig 5E). These findings lead us to speculate that TRIM32 resides in the endosome and may impair cytosolic entry of VEEV after endosomal membrane fusion prior to genome translation.

To test this model, we investigated the fate of incoming VEEV nucleocapsids using VEEV-NLuc/Cap (Fig 6A and 6B). We found that the incoming NLuc/Cap signal rapidly decreases starting at 30 minutes post entry, suggesting that nucleocapsids may undergo degradation or structure disruption in the presence of TRIM32 (Fig 6C). A similar phenomenon has been demonstrated during Dengue virus [46] and Yellow fever virus [47] infection, with degradation of incoming capsids being important for initiation of viral genome translation. Next, the stability of the NLuc/Cap of VEEV was determined in HeLa-TRIM32 or HeLa-Fluc cells by VEEV-NLuc/Cap infection in the presence or absence of cycloheximide. We found that NLuc/Cap is less stable in TRIM32 expressing cells compared to HeLa-Fluc cells (Fig 6D and 6E). Our replicon assay showed that TRIM32 does not affect transfected viral RNA-mediated translation and replication (Fig 3C and 3D). Thus, we hypothesized that TRIM32 may destabilize the incoming nucleocapsids resulting in inhibition of primary genome translation. To verify this hypothesis, we engineered a reporter VEEV that has an NLuc fusion at the C-terminus of nonstructural protein 3, hereafter termed VEEV-nsP3/NLuc (S4 Fig). As shown in Fig S4, HeLa-TRIM32 and HeLa-Fluc cells were infected with VEEV-nsP3/NLuc at MOI of 1, and cells were collected at multiple time points corresponding to steps of initial viral infection. Compared to HeLa-Fluc, the initial translation of VEEV-nsP3/NLuc, as indicated by NLuc activity, is significantly lower in HeLa-TRIM32 cells (Fig 6F). To focus on the effects of TRIM32 on viral translation, ML336, a VEEV nsP2 specific inhibitor [48] was added to block viral genome replication-mediated amplification of NLuc signal. Again, the NLuc activity was lower in HeLa-TRIM32 cells during at all time points tested (Fig 6G). Together, by using sensitive viral tools, our data demonstrate that TRIM32 impairs a late entry step in a manner that leads to impaired translation of encapsidated viral RNA.

**Fig 6.**
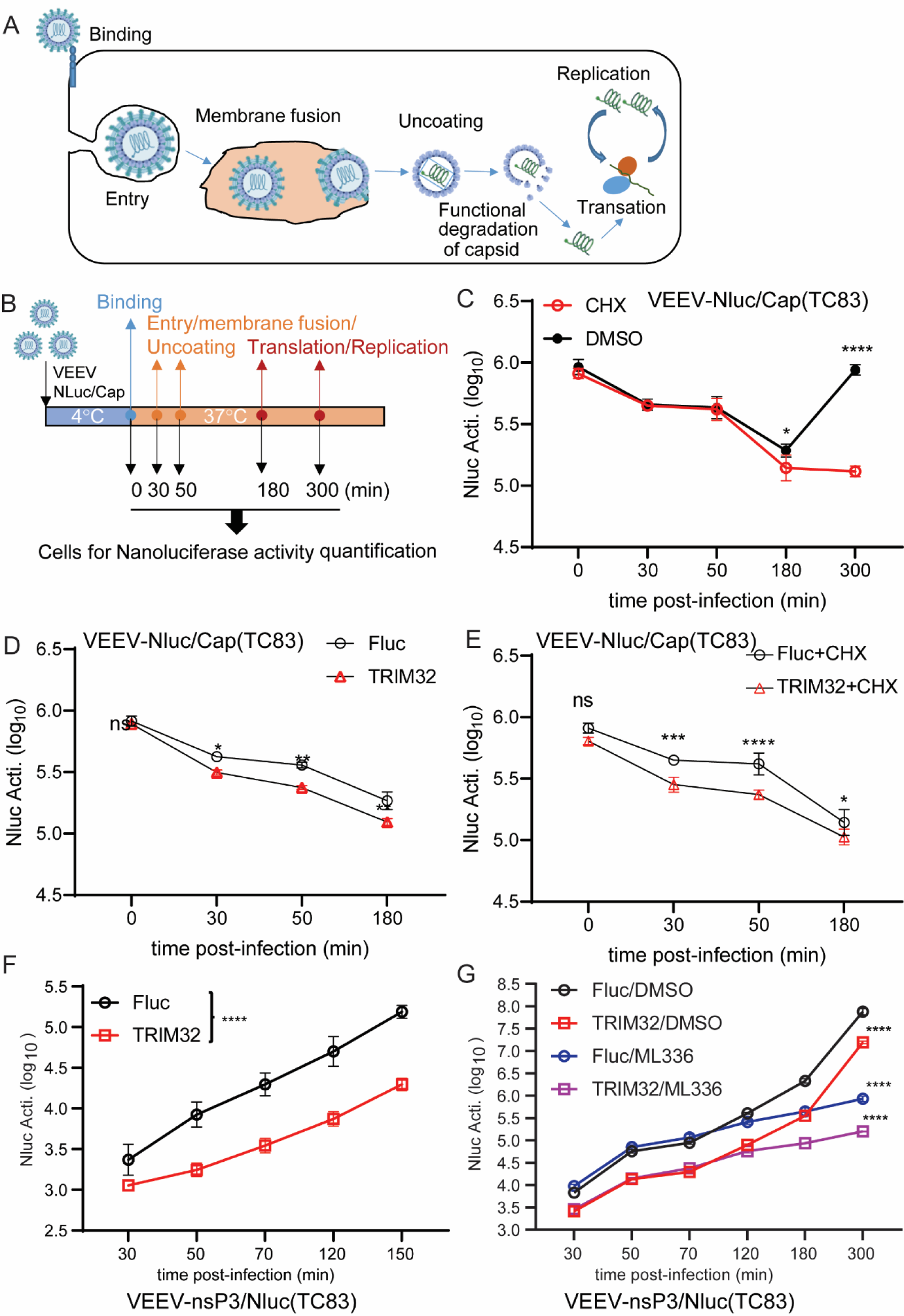
TRIM32 impairs the translation of incoming viral genome. **A.** Schematic of the life cycle of VEEV. **B.** Schematic illustration of stability assay of VEEV nucleocapsids by using VEEV-NLuc/Cap infection. **C**. HeLa cells were infected VEEV-NLuc/Cap at 4°C for 1.5h, then the cells were washed and collected at the indicated time post temperature shift. **D.** HeLa-Fluc or HeLa-TRIM32 cells were infected with VEEV-NLuc/Cap at 4°C for 1.5h, then the cells were washed and collected at the indicated time post temperature shift. **E.** HeLa-Fluc or HeLa-TRIM32 cells were infected VEEV-NLuc/Cap at 4°C for 1.5h, then the cells were washed and collected at the indicated time post temperature shift in the presence or absence of CHX (75 ug/mL). **F**. HeLa-Fluc or HeLa-TRIM32 cells were infected with 1 MOI of VEEV-nsp3/NLuc at 37C for 20min, then the cells were washed and replaced with fresh media. The cells were collected at the indicated time points post infection, and the Nanoluc activity was measured. **G**. HeLa-Fluc or HeLa-TRIM32 cells were infected with 1 MOI of VEEV-nsp3/NLuc at 37°C for 20min, then the cells were washed and replaced with fresh media containing DMSO or 1μM ML336. The cells were collected at the indicated time points post infection, and the Nanoluc activity was measured. Data represent averages of independent biological replicates and are presented as means ± SD (n=3). Statistical significance was determined by 2-way ANOVA with Sidak’s multiple comparisons test (****P<0.0001).

### TRIM32 R394H and D487N pathogenic mutants fails to restrict VEEV infection

Previous studies report that mutations in the TRIM32 NHL domain are highly associated with Limb-girdle muscular dystrophy H2 (LGMDH2) or R8 (LGMDR8), which cause autosomal recessive muscle disease [29][49]. However, whether these pathogenic mutations influence the antiviral activity of TRIM32 has not been demonstrated. Here, we established HeLa stable cells expressing four TRIM32 mutants, including C39/44A that inactivates E3 ubiquitin ligase activity, P130S, which is associated with Bardet-Biedl syndrome (BBS11), and two well-characterized mutants R394H and D487N that are associated with Limb-girdle muscular dystrophy (Fig 7A). Cells expressing TRIM32 mutants were infected with VEEV-TC83-GFP, and virus infectivity was quantified by flow cytometry. We found that the P130S mutant retains antiviral activity against VEEV, but C39/44A, R394H, and D487N failed to inhibit VEEV infection (Fig 7B). Western blot analysis showed the expression of wild type and mutated TRIM32 proteins were not identical, as cells expressing wild-type or P130S TRIM32, but not C39/44A, R394H, or D487N possess a higher molecular weight band (Fig 7B). This higher molecular weight band has been characterized as monoubiquitinated TRIM32 [50], suggesting that loss of TRIM32 antiviral function correlates with a loss of TRIM32 monoubiquitination. Next, we tested whether the intracellular distributions of C39/44A, R394H, and D487N were altered compared to wild-type TRIM32. Indeed, the confocal images show that P130S behaved like the wild type TRIM32, which both form cytoplasmic bodies of similar size. However, C39/44A mutant TRIM32 appeared to form massive aggregates, while both R394H and D487N failed to form cytoplasmic bodies and showed diffuse cytoplasmic distribution (Fig 7C).

**Fig 7.**
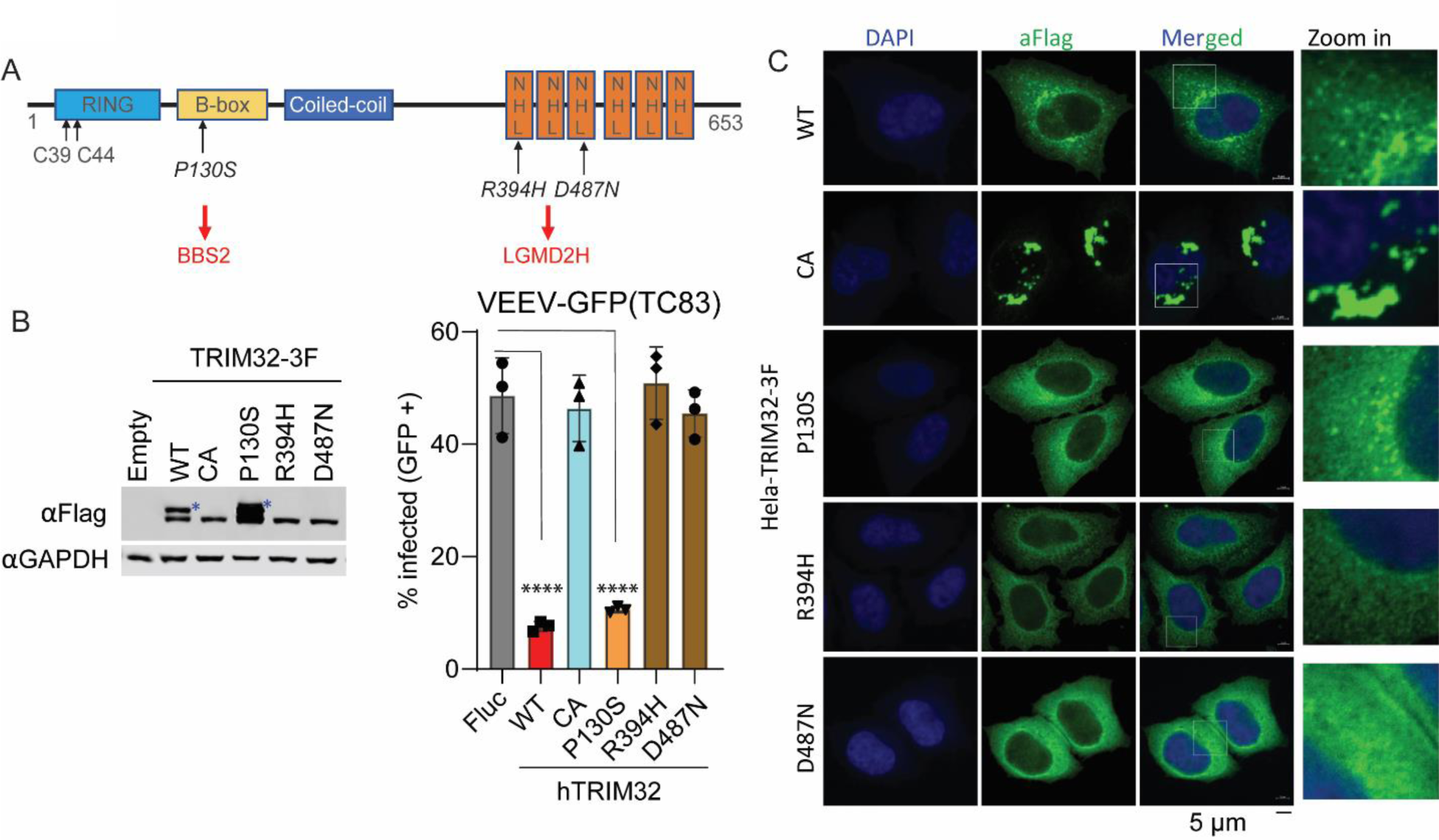
TRIM32 LGMD2H mutants do not exhibit antiviral activity against VEEV. **A.** Schematic of TRIM32 protein with domain features and important residues. **B.** Left: Western blot analysis of expression of wild type and mutated TRIM32 in HeLa stable cells. Right: VEEV-TC83-GFP infectivity in the indicated HeLa stable cells. The cells were infected VEEV-TC83-GFP at 0.1 MOI for 24h, viral infectivity was measured by flow cytometry. The asterisks denote the monoubiquitinated TRIM32. **C.** Microscope analysis the intracellular distribution of wild type and the indicated TRIM32 mutants in HeLa cells. Nuclei are stained with DAPI, blue signal. Wild type and mutated TRIM32, green signal. Scale bar, 5 μM. Data represent averages of independent biological replicates and are presented as means ± SD (n=3). Statistical significance was determined by unpaired students’ t-test for B (****P<0.0001).

Collectively, our data demonstrate that the LGMDH2 associated TRIM32 mutants fail to inhibit VEEV infection, while BBS11 related mutant P130S retains antiviral activity. Moreover, E3 ubiquitin ligase activity of TRIM32 is required for inhibition of VEEV.

## Discussion

In this study, we screened a library of E3 ligases and identified TRIM32 as a restriction factor against alphaviruses, including VEEV, SINV, and ONNV (Fig 1, S1). We also show that TRIM33 expression, a negative regulator of host innate immune response [52,53], significantly promoted VEEV-TC83 infection (Fig 1A and 1B). TRIM33 can also act as direct restriction factor that suppresses HIV-1 replication by targeting the viral integrase for proteasomal degradation [54]. Thus, the potential mechanism of TRIM33-mediated enhancement of VEEV warrants further study.

TRIM32 belongs to C-VII subclass membrane of the TRIM protein family, which contains six NHL (NCL-1, HT2A, LIN-41) repeats in its C-terminus [55]. Human TRIM32 is expressed in many tissues, including skeletal muscle, brain, and heart [29]. As a E3 ubiquitin ligase, TRIM32 plays important roles in several cellular process, including differentiation, tumor suppression, regeneration, microRNA process, muscle physiology, and antiviral innate immunity [30,34,35,37]. TRIM32 can also positively regulate host antiviral innate immunity by targeting STING for K63-linked ubiquitination [34], which results in a broad-spectrum of antiviral effects. In addition, TRIM32 can also be an intrinsic restriction factor. During IAV infection, basally expressed TRIM32 targets PB1 for proteasomal degradation, which inhibits infection prior to interferon-mediated effects [37].

In this study, we expand the role of TRIM32 as a restriction factor by demonstrating that it inhibits alphaviruses independently of its known role in regulating STING-IFN pathways (Fig 1, 2). Mechanistically, we found that TRIM32 does not affect the viral binding, internalization of virions, or membrane fusion (Fig 4). Our data also suggest TRIM32 does not inhibit primary translation of viral genome replication. These conclusions were reached with several tools. Using the replication inhibitor ML366, we found that ML366 exerted no effects on VEEVN-nsp3/NLuc during the first 120 minutes of infection, whereas TRIM32 suppressed nsp3/NLuc activity at the earliest time point of 30 minutes (Fig 6G). ML366 began suppressing nsp3/NLuc after 120 minutes, indicating that replication begins around this time. TRIM32 did not suppress this early replication phase burst in nsp3/Nluc activity, suggesting that viral genome amplification from the RNA-dependent RNA polymerase was likely not affected. Additional support for a lack of effect on replication come from entry bypass experiments, in which TRIM32 did not suppress replication of transfected full length viral RNAs or replicons (Fig 3A, B). We additionally infer from our data that TRIM32 does not directly suppress primary translation of incoming RNAs. The VEEV-Rluc replicon activity was monitored from 1h to 6h, and TRIM32 had no effect at any time point, including 1 hour (h), suggesting that translation if the incoming RNA is unaffected (Fig 3 C, D). We note limitations of this interpretation below.

Our studies suggest TRIM32 inhibits a late entry step, likely after membrane fusion but prior to translation of viral RNA. We speculate TRIM32 may affect late entry processes related to uncoating based on studies with a new viral tool, VEEV-NLuc/Cap, which has NLuc fused to capsid in the virion (Fig S2). TRIM32 expression resulted in decreased NLuc/Cap reporter activity at very early time points (30 min) after cold binding the virus and allowing internalization with a temperature shift (Fig 6D). This effect also occurred in the presence of cycloheximide, indicating that protein synthesis is not needed for TRIM32-dependent inhibition. Taken together, these replication cycle studies point to a TRIM32-mediated inhibition of VEEV at a late step in the viral entry pathway. This conclusion is consistent with proximity labeling proteomics and microscopy studies indicating that TRIM32 resides in endosomes (Fig 5), which are important for cytosolic entry of VEEV [38].

We additionally examined TRIM32 mutations associated with Bardet-Biedl syndrome (BBS11) and limb-girdle muscular dystrophies (LGMDs) affect anti-alphavirus activity [29,32]. We found that P130S behaves similar to wild type TRIM32 with respect to viral inhibition and monoubiquitination, but R394H and D487N lost antiviral activity against VEEV-TC83 and did not express the larger second band, suggesting loss of monoubiquitination (Fig 7B). These defects also correlated with altered intracellular distributions (Fig 7C). We also show that E3 ligase activity is required for TRIM32-mediated inhibition of VEEV. However, we could not assess whether UPS inhibitors can subvert TRIM32-mediated inhibition of VEEV, because the host UPS also required for virus infection [19,21]. Previous study reported that TRIM32 mutants R394H and D487N lost monoubiquitination feature, but still remain the ability to self-associate. We also show that the monoubiquitination of TRIM32 is likely critical for its antiviral activity, as mutants lacking monoubiquitination also lack anti-VEEV function.

Summarily, our findings implicate TRIM32 as novel host factor that inhibits alphaviruses. Our data suggest that TRIM32 inhibits a late step in viral entry, possibly related to capsid uncoating. However, our study has several limitations that, if addressed in future studies, would help further refine the biological relevance of TRIM32 during alphavirus infection and the current mechanistic model. 1) Additional TRIM32 gene silencing studies in primary cells will help confirm our studies in immortalized cells that TRIM32 functions as a cell intrinsic restriction factor. 2) TRIM32 does not appear to be an ISG [62], emphasizing the need for additional studies to understand how TRIM32 expression is regulated. 3) Our mechanistic studies did not include assays to directly investigate endosomal fusion, and R18-labeled virus could assist in further narrowing the proposed late entry mechanism. 4) While mutagenesis indicates that the E3 ligase activity of TRIM32 is important for its effects on VEEV, additional studies are needed to identify whether a viral or host factor is ubiquitinated by TRIM32, and what the consequence of this modification is. 5) The association of TRIM32 mutations with certain human diseases raises the question of whether individuals bearing these SNPs in TRIM32 are at higher risk for alphavirus infection. Additional studies in human populations or genetically defined animal models could help address this question.

## Materials and methods

### Cell culture and viruses

293T, HeLa, Huh7.5, U2OS, C2C12, and BHK-21 cells were maintained in Dulbecco’s Modified Eagle Medium (DMEM, Gibco) supplemented with 10% FBS (Gibco), 1% Penicillin-streptomycin (Gibco), and 1× non-essential amino acids (NEAA; Gibco). All these cells were cultured at 37°C with 5% CO2. The viruses that used in this study were propagated as previously described: VEEV-GFP [63], ONNV-GFP, SINV-GFP, and CVB3-GFP [41]. ZIKV-GFP [64].

### Plasmids and molecular cloning

Lentiviral-based expression vector pTRIP.CMV.IVSb.ires.TagRFP-DEST [63], pSCRBBL-DEST, or pSCRPSY-DEST [65] were used for expression of genes of interest and generated using Gateway cloning.

To generate TRIM32/mCherry or TRIM32mNeoGreen expressing constructs, overlapping PCR was performed to fuse mCherry or mNeonGreen to the C-terminus of TRIM32. TRIM32 C39/44A, P130S, R394H, and D487N mutants were generated by overlapping PCR.

CRISPR–Cas9 cloning and knockout bulk cells were prepared as previously described. Oligos encoding sgRNAs for generating knockout cells using CRISPR–Cas9 were cloned into the lentiCRISPRv2 plasmid (a gift from F. Zhang, Addgene plasmid 52961) as previously described [66]. LentiCRISPRv2 clones containing guide sequences were sequenced, purified, and used for lentiviral production as described above. For generating bulk knockout cells, HeLa or C2C12 cells were transduced with the lentiCRISPRv2-derived lentivirus for 48 h, then reseeded into complete DMEM containing 4 µg/ml puromycin for 3 days of selection to kill untransduced cells.

### Lentivirus production, transduction, and viral infection assays

Lentiviral production and transduction was performed as previous described [63]. Briefly, 8 × 10^5^ 293T cells in 6-well plates or 3 × 10^5^ cells in 24-well plates were co-transfected with plasmids expressing E3 (pTRIP-E3 or pSCRPSY-E3) [39], HIV-1 gag–pol and VSV-G in a ratio of 1/0.8/0.2, respectively. For 6-well plate transfection, 6 μl X-tremeGENE HP (Roche) was combined with 2.0 μg total DNA in 100 μl Opti-MEM (Gibco). For 24-well plate transfection, 1.5 μl FuGENE (Roche) was combined with 0.5 μg total DNA in 25 μl Opti-MEM. Transfections were carried out for 6 h, followed by a medium change to DMEM containing 3% FBS. Supernatants were collected at 48 h and 72 h, pooled, cleared by centrifugation and stored at −80 °C. For lentiviral transduction, HeLa cells were seeded in to 24-well plates at a density of 2 × 10^4^ cells per well and transduced with lentivirus in transduction medium that containing 3% FBS, 20 mM HEPES, and 4 μg/mL polybrene.

### E3 screen

HeLa cells were seeded at a density of 2×10^4^ cells per well in 24-well plates and cells were then transduced with E3 lentiviruses in transduction medium. Cells were split and replated into 24-well plate at a ratio of 1:4 at 48 h post-transduction. 16-24 h later, the cells were infected with the VEEV-TC83-GFP at 0.1 multiplicity of infection (MOI. 1 MOI = 1 plaque formation unit per cell). The cells were harvested at 24 h post-infection for FACS-based infectivity analysis as previous study described [63].

### In vitro transcription of viral infectious RNA and replicon RNA

VEEV-TC83 viral RNA and replicon RNA were in vitro transcribed using mMessage mMachine SP6 Transcription kit (Thermo Fisher, Cat#AM1340). RNA was purified from the transcription reaction using RNeasy mini kit (Qiagen, Cat#74106) and quantified by Nanodrop. Transfection of viral infectious RNA and replicon RNA was performed with the TransIT®-mRNA Transfection Kit (Mirus Bio, Cat#MIR2250). To produce virus from in vitro transcribed RNA, BHK-21 cells were seeded in a 100cm dish at a density of 2 x 10^6^ cells the day before transfection. A total of 4 µg of viral RNA was then transfected in each dish, and virus was harvested 36 to 48 hrs post-transfection. For replicon assays, cells were plated at a density of 5 x 10^4^ cells per well in 24-well plates the day before transfection. 100 ng viral replicon RNA was then transfected into cells. Transfected cells were then harvested at the indicated time points using Renilla lysis buffer. Renilla luciferase activity was quantified using the Renilla Luciferase Assay System (Promega, Cat# E2820).

### Viral binding and entry assay

Cells were seeded in 24-well plates at a density of 1 x 10^5^ cells per well the day before infection. The cells were infected with VEEV-TC83-GFP at MOI of 20 and incubated with cells for 1.5 h at 4°C with horizontal shaking every 20-30 minutes. Unbound viruses were then washed off using chilled PBS. After the final wash, 200 µL FBS free DMEM was added into the wells, and the plates were placed in −80°C for at least 2h. After three freeze-thaw cycles, bound and internalized virus was quantified by real-time PCR detecting VEEV-TC83 nsP3 gene.

For VEEV-NLuc/Cap binding assay, cells were seeded in 24-well plates at a density of 5 x 10^4^ cells per well the day before infection. The cells were infected with VEEV-NLuc/Cap at MOI of 1 or 10 and incubated with cells for 1.5 h at 4°C with horizontal shaking every 20-30 minutes. Unbound viruses were then washed off using chilled PBS for 4 washes. After the final wash, add 150 µL NP40 lysis buffer into the wells and plates were stored in the −80°C freezer. The NLuc activity was determined by using Nano-Glo Luciferase Assay System.

### Plaque assay

BHK-21 cells were seeded in 12-well plates at a density of 2 x 10^5^ cells per well the day before infection. The cells were infected with diluted virus in 200µL medium and bound to cells at 37°C for 1.5h. Then the cells were covered by Avicel overlay medium [67], and incubated at 37°C for 3 days. Plaques were then visualized by crystal violet staining.

### Real-time PCR

Total RNA from tissue culture cells was extracted using RNeasy mini kit (Qiagen, Cat#74106) following the manufacturer’s instructions. Total RNA from mouse tissue was extracted using TRIzol (Invitrogen). For RT-PCR assays, SuperScript IV First-strand synthesis system (Invitrogen, Cat#18091050) was used to generate the complementary DNA template.

### Immunofluorescence assay and confocal microscopy

For confocal immunofluorescence studies, the indicated antibodies and dyes were diluted in 5% BSA in PBS as below: mouse anti-Flag (ProteinTech, Cat#66008-4-Ig, 1:5,000), goat-anti-mouse Alexa Fluor-488 (Invitrogen, 1:1,000), goat-anti-rabbit Alexa Fluor-488 (Invitrogen, 1:1,000), goat-anti-rabbit Alexa Fluor-647 (Invitrogen, 1:1,000), DAPI (4’,6-diamidino-2-phenylindole) was used for staining of nuclei. Images were collected by using ZEISS LSM 880 with Airyscan system at the core facility at Life Science Institute, Zhejiang University.

### Protein lysate preparation and Western blot

Tissue culture cells were lysed in Nonidet P-40 (NP-40) buffer (50 mM Tris pH 7.4, 150 mM NaCl, 1% (v/v) NP-40, 1 mM EDTA, protease inhibitor cocktail) and incubated on ice for 30min, followed by centrifugation at 13,000 rpm for 20 min at 4°C. For mouse tissue lysates, fresh tissues were collected in 2ml round-bottom Eppendorf tubes containing NP40 lysis buffer and cut into small pieces using scissors. Western blot analyses were then performed as previously described [68].

### Proximity labeling proteomics

HeLa cells were transduced with lentivirus expressing TurboID.HA or TRIM32.TurboID.HA to generate HeLa-TurboID.HA and HeLa-TRIM32.TurboID.HA. Cells were seeded in 100mm dishes at a density of 5 x 10^6^ cells per dish. The cells were mock infected or infected with VEEV-TC83-GFP at MOI of 25 in the presence of biotin at a concentration of 500 μM. The cells were collected at 6 hour post infection (hpi), and biotinylated proteins were enriched as previously described [69]. The biotinylation efficiency confirmed by western blot. To prepare proteomic samples for mass spectrometry analysis, the enriched proteins were separated on a gradient gel (GenScript, 4–20% precast polyacrylamide gel) at 120V for 5∼10 min, until the protein ladder ran ∼1cm into gel. Then the gel was stained with Coomassie blue (G-250) for 1h at room temperature. After destaining in buffer containing 45%(v/v) methanol, 10% (v/v) acetic acid in HPLC-grade water the stained area was cut into slices and each slice was placed into an Eppendorf 1.5 mL tube, which had been previously rinsed with 50% acetone and millipure water. Proteins were identified by mass spectrometry by the Protein Chemistry Facility at the Center for Biomedical Analysis of Tsinghua University.

### Antibodies and reagents

The following antibodies were used for immunoblotting: mouse anti-Flag (Sigma, M2, 1:5,000), rabbit anti-Flag (ProteinTech, 1:5,000), mouse anti-GAPDH (Abclonal, Cat#AC002, 1:5,000), rabbit anti-TRIM32 (ProteinTech, Cat# 10326-1-AP, 1:1,000), rabbit anti-STING (ProteinTech, Cat#19851-1-AP, 1:1,000), mouse anti-STAT1 (Proteintech, Cat#66545-1-Ig, 1:1,000), Rabbit-anti-TBK1 (ProteinTech, Cat# 28397-1-AP, 1:1,000). For western blotting, antibodies were diluted in 5% (w/v) non-fat dry milk in TBST buffer. For immunofluorescence assay, the antibodies were diluted in 5% (w/v) BSA in PBS. D-biotin (Sangon Biotech, Cat# A100340), Streptavidin MagBeads (GenScript, Cat#L00936), Nano-Glo Luciferase Assay System (Promega, Cat#N1110), Renilla-Lumi Plus (Beyotime, Cat#RG066S), Cycloheximide (4A Biotech, Cat#FXP130-10), ML336 (TargetMol, Cat#T21900-1).

## Supporting information

**S1 Fig.** Knockout of STING in U_2OS-TRIM32 or U-2OS-Fluc cells. Cells were infected with the indicated viruses and virus infectivity was quantified by flow cytometry.

**S2 Fig.** Schematic illustration of VEEV-NLuc/Cap and viral protein analysis. A. Illustration of VEEV-NLuc/Cap. B. VEEV-TC83 and VEEV-NLuc/Cap were purified by ultracentrifugation, ponceau S staining. Western blot analysis using antibody against viral capsid protein was employed to analyze viral proteins from virions.

**S3 Fig.** Proximity labeling proteomics approach to identify TRIM32 interacting proteins. A. Western blot analysis of TRIM32-TurboID system. HeLa-TRIM32.TurboID.HA or HeLa-TurboID.HA cells were mock infected or infected with VEEV-TC83-GFP at MOI of 25 for 6 hrs in the presence of biotin at a concentration of 500μM. The biotinylated proteins were purified by using Streptavidin MagBeads. B. The enriched proteins in each groups were analyzed by using Venny 2.1 (Venny 2.1.0 (csic.es)).

**S4 Fig.** Schematic illustration of VEEV-nsP3/NLuc and related workflow. SGp: subgenomic promoter, NLuc, Nanoluciferase.

S Table DNA oligos used in this study.

## Acknowledgements

We kindly thank the Protein Chemistry Facility at the Center for Biomedical Analysis of Tsinghua University for mass spectrometry analysis. We also thank the imaging core facility at Life Sciences Institute, Zhejiang University.

## Data Availability Statement

All relevant data are within the paper and its Supporting Information files.

## Author Contributions

**Conceptualization:** John W. Schoggins, Wenchun Fan.

**Formal analysis:** Yifan Xie, Lingdong Xu, Wenchun Fan.

**Funding acquisition:** John W. Schoggins, Wenchun Fan.

**Investigation:** Yifan Xie, Jie Cao, Shuyi Gan, Lingdong Xu, Dongjie Zhang, Suhong Qian, Wenchun Fan.

**Methodology:** Yifan Xie, Jie Cao, Shuyi Gan, Lingdong Xu, Dongjie Zhang, Suhong Qian, Qiang Ding, Feng Xu, John W. Schoggins, Wenchun Fan.

**Project administration:** John W. Schoggins, Wenchun Fan.

**Resources:** Qiang Ding, John W. Schoggins, Wenchun Fan.

**Supervision:** John W. Schoggins, Wenchun Fan.

**Visualization:** Yifan Xie, Jie Cao, Shuyi Gan, Lingdong Xu, Wenchun Fan.

**Writing – original draft:** Wenchun Fan.

**Writing – review & editing:** Yifan Xie, Jie Cao, Shuyi Gan, Lingdong Xu, Qiang Ding, John W. Schoggins, Wenchun Fan.

---

**S1 Fig.**
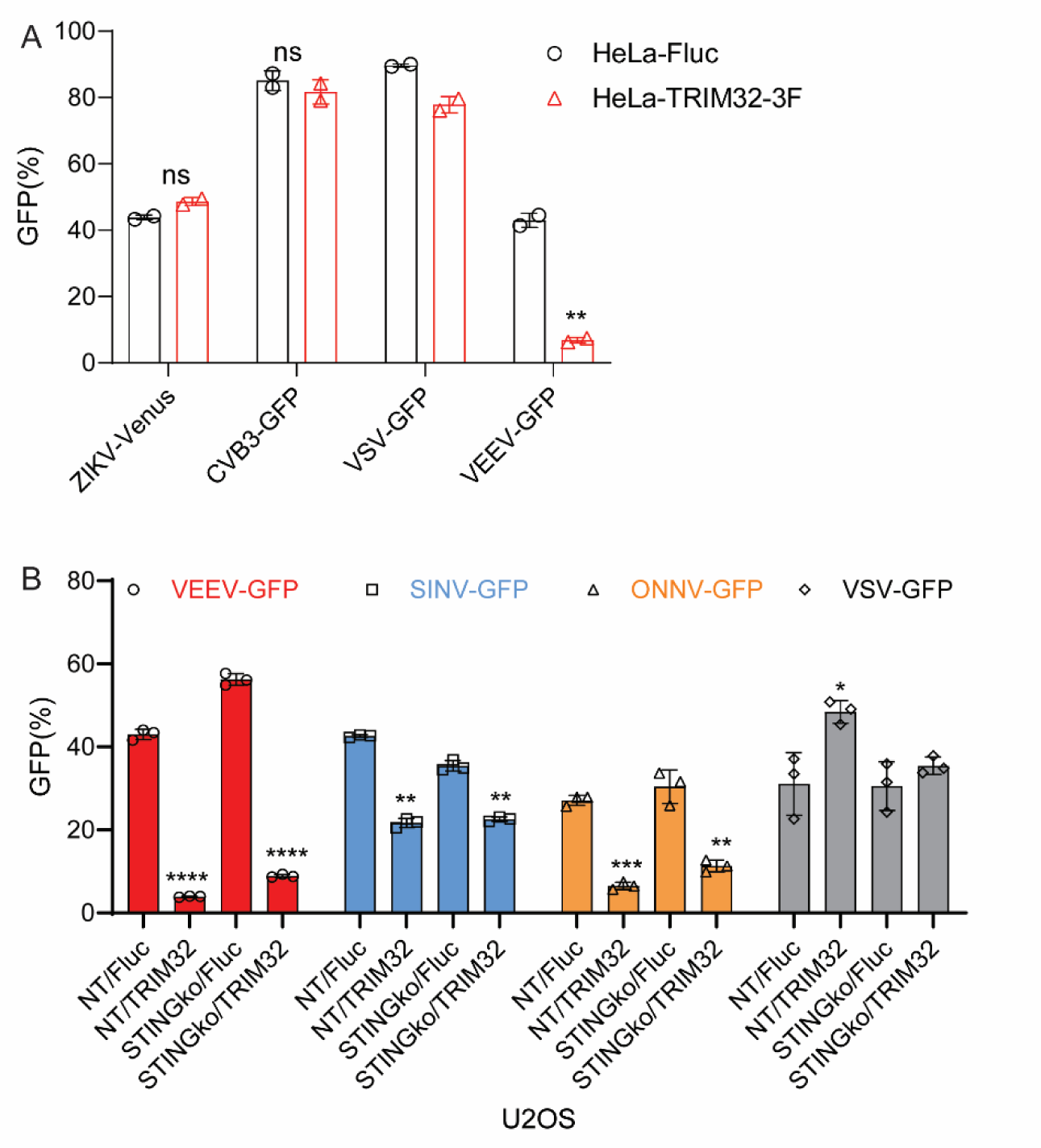
A. HeLa-Fluc or HeLa-TRIM32-3F were infected with the indicated viruses, and viral infectivity was quantified by flow cytometry. B. STING was silenced in U2-OS-TRIM32 or U2-OS-Fluc cells. Cells were infected with the indicated viruses, and virus infectivity was quantified by flow cytometry. Statistical significance was determined by unpaired students’ t-test for A and B (*P<0.05,**P<0.01, ***P<0.001, ****P<0.0001). ns, no significant..

**S2 Fig.**
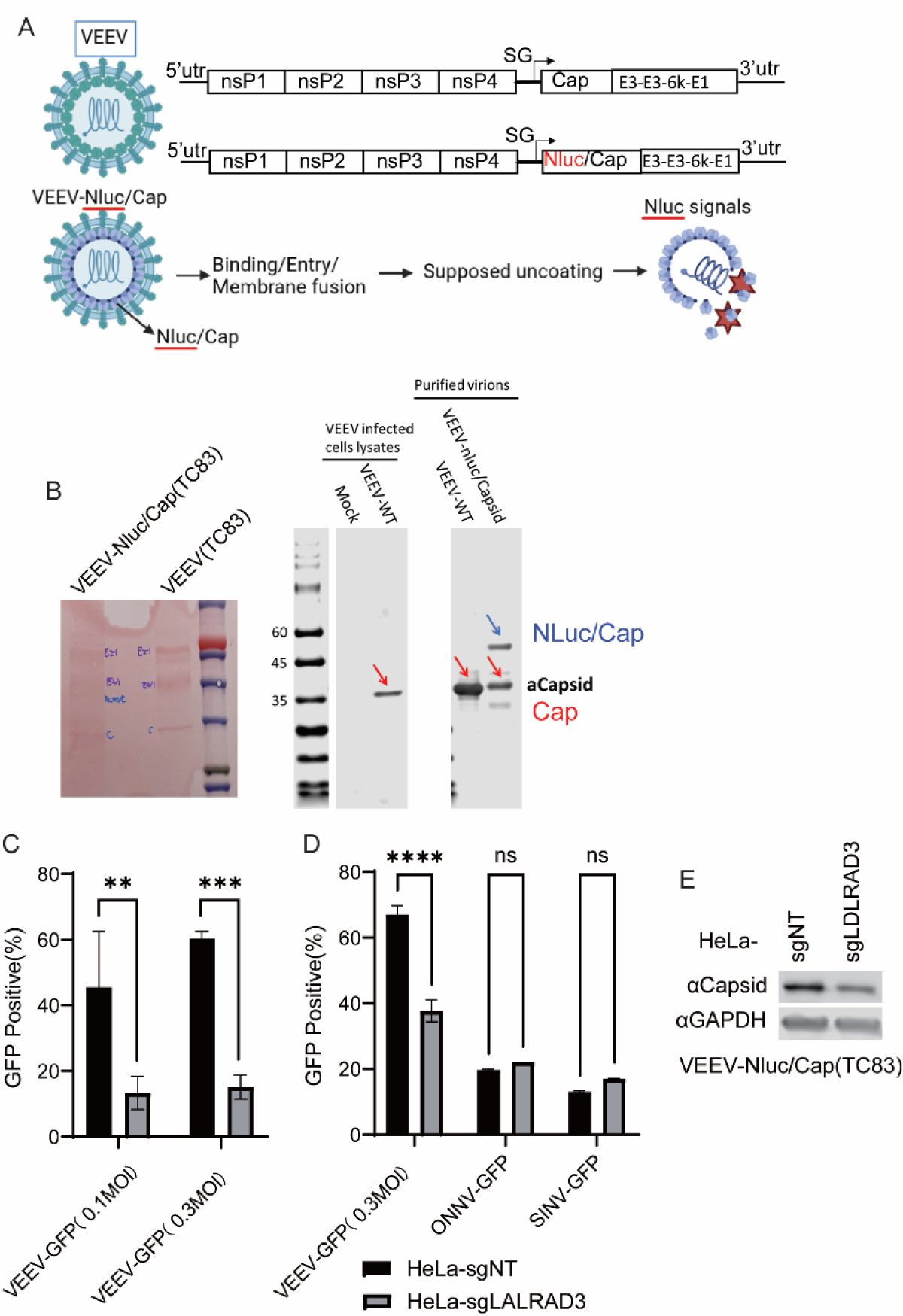
A. Illustration of VEEV-NLuc/Cap. B. VEEV-TC83 and VEEV-NLuc/Cap was purified by ultracentrifugation, and virion proteins were visualized by ponceau S staining and Western blot analysis using antibody against viral capsid protein. C. HeLa cells stable expressing dual-sgRNA targeting VEEV receptor LDLRAD3 or non-specific targeting sgRNA were infected VEEV-TC83-GFP at the indicated MOI for 24h, and virus infectivity was quantified by flow cytometry. D. HeLa cells stable expressing dual-sgRNA targeting VEEV receptor LDLRAD3 or non-specific targeting sgRNA were infected VEEV-TC83-GFP, SINV-GFP, and ONNV-GFP at MOI of 0.1 for 24h, and virus infectivity was quantified by flow cytometry. Statistical significance was determined by unpaired students’ t-test for C and D (**P<0.01, ***P<0.001, ****P<0.0001). ns, no significant.

**S3 Fig.**
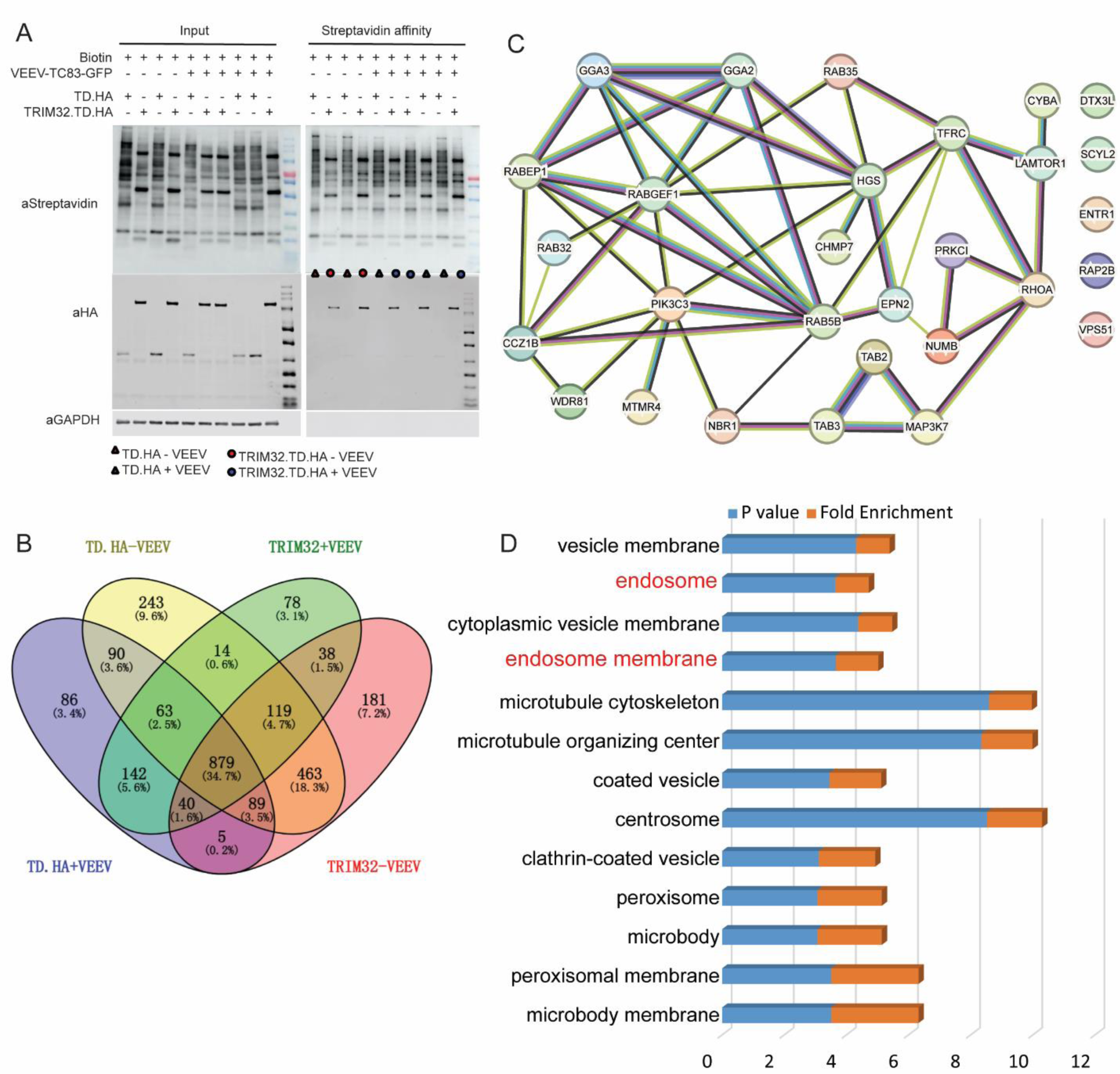
Proximity labeling proteomics approach to identify TRIM32 interacting proteins. A. Western blot analysis of TRIM32-TurboID system. HeLa-TRIM32.TurboID.HA or HeLa-TurboID.HA cells were infected with or without VEEV-TC83-GFP at MOI of 25 for 6 hrs in the presence of biotin at a concentration of 500μM. The biotinylated proteins were purified by using Streptavidin MagBeads. B. The enriched proteins in each group were analyzed by using Venny 2.1 (Venny 2.1.0 (csic.es). **C.** The cellular component analysis of TRIM32-enriched proteins by using PANTHER [70]. **D.** The network of TRIM32 proximity labeled endosome associated proteins.

**S4 Fig.**
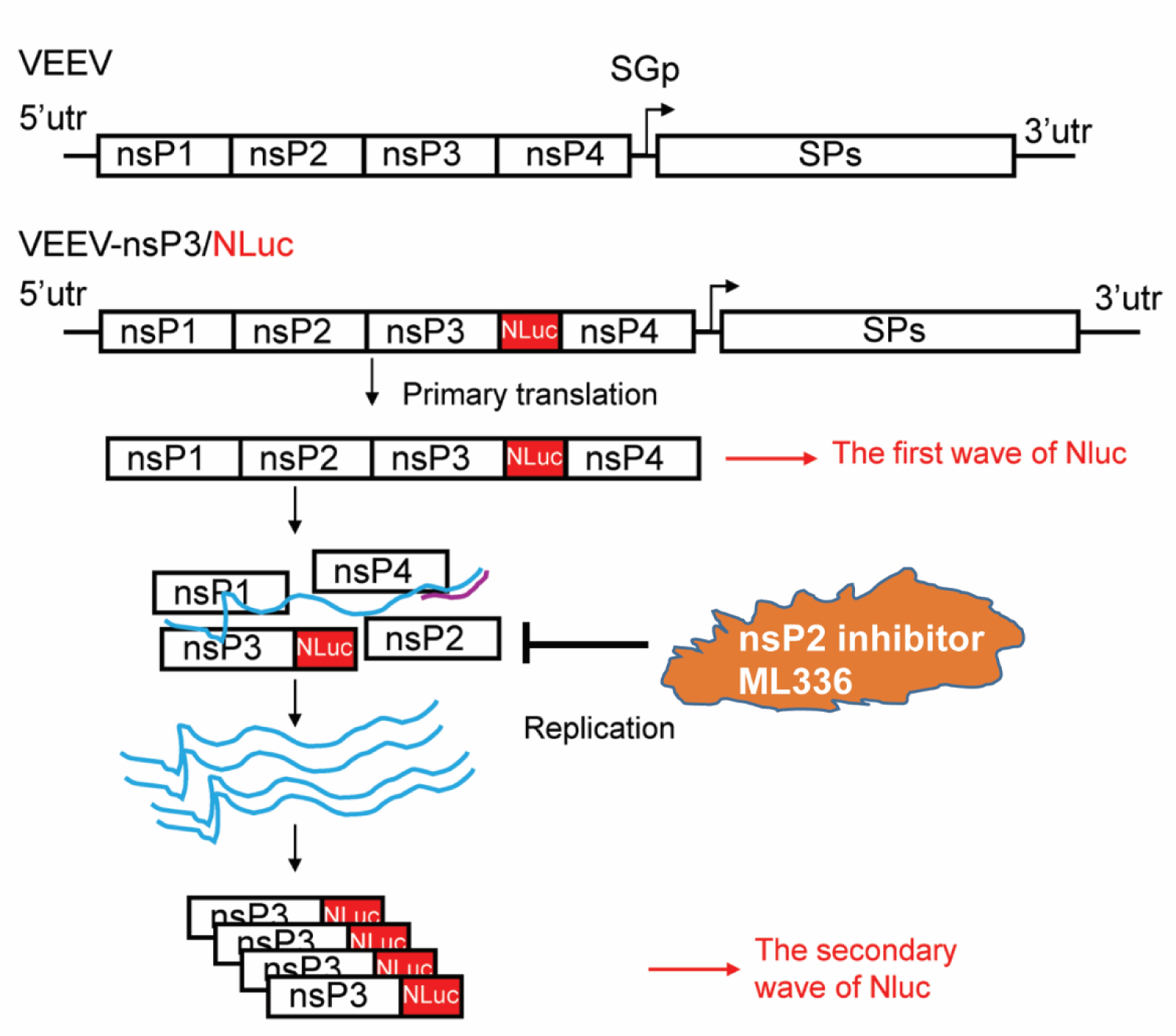
Schematic of VEEV-nsP3/NLuc and related workflow. SGp: subgenomic promoter, NLuc, Nanoluciferase.

